# A sensitive and specific microchimerism detection method using custom designed genotyping and companion qPCR assays

**DOI:** 10.64898/2026.10.22.761108

**Authors:** John Houck, Neta Simon, Blair Armistead, Alisa Kachikis, Heather B. Jaspan, Lisa Frenkel, Swati Shree, Whitney E. Harrington

## Abstract

Since first reports of the presence of a small number of cells from one individual persisting in another, researchers have developed a number of methods to detect microchimerism. Molecular based detection platforms range from gel electrophoresis to quantitative PCR to Next-Gen Sequencing. Central to all approaches is the need for informative polymorphic loci to effectively distinguish microchimeric cells from the host. However, we observed in our studies of maternal microchimerism that currently published approaches can limit the number of samples available for assessment due to the inability to identify an informative marker of the donor in the recipient. This was most striking in our studies of populations of African descent in which the use of HLA-based markers, a common approach, prevented the testing of 30-50% of participants. Other potential options were cost prohibitive or lacked sufficient sensitivity. To address this challenge, we developed a novel, simple, and low-cost approach that leverages global population frequency databases for polymorphism selection, reduces cost by amplifying multiple targets in one multiplex assay to complete genotyping, and has high sensitivity and specificity using rigorously designed qPCR assays to quantify microchimerism. Herein, we outline our design process, detailed assay protocol, and assessment of MMc using this approach in two different populations of infants from the United States and South Africa.

## INTRODUCTION

A variety of molecular methods have been developed to detect chimerism and/or microchimerism in samples for clinical and research purposes. Methodologies using short tandem repeats (STR), single nucleotide polymorphisms (SNPs), HLA discordance, and Y chromosome detection have all been reported but can have limitations either in sensitivity, substrate requirements, or cost (1, 2). For example, limit of chimerism detection using STR has been reported at 1-5% (3–5) and using custom qPCR at 0.1% (6). Other more sensitive assays such as the qPCR based QTRACE® Analysis System (JETA Molecular, Utrecht, Netherlands) and the KMRtype Genotyping Kit (GenDx Products, San Diego, USA) use individual qPCR assays as screening tools to identify informative markers, thus requiring enough genomic DNA (gDNA) to both screen and test in up to 40 assays (7, 8). Other available assays using next generation sequencing (NGS) have been reported including the One Lambda Devyser Chimerism NGS Assay (Thermo Fisher Scientific, Los Angeles, USA) and ScisGo Chimerism Multi-Donor Assay (Scisco Genetics, Inc.) but can have variable sensitivity (0.2-1%) due to the limitations of the NGS platform (9–15). Other novel, custom-designed NGS assays allow for potential interrogation of a wide number of polymorphisms in one assay; however, the throughput and cost make it challenging to explore population based microchimerism using this approach (16). Further, many of the available assays are designed for clinical testing; as such, the per sample cost tends to be higher than many research budgets allow. A low-cost approach that allows for assessment of a large number of samples but still retains sensitivity and specificity is currently lacking.

Common to all current chimerism assays is reliance on polymorphisms to identify the donor cells in the background of the recipient, making target polymorphism selection key to the success of the approach. Custom assays can be limited when studying ethnic groups different from those used for initial assay design, as polymorphism allele frequency can vary substantially between population groups. We observed this limitation firsthand when utilizing published assays targeting non-shared non-inherited HLA polymorphisms to detect maternal microchimerism (**MMc**) (17, 18). For example, we observed that more than a third of samples in a Malian cohort (*unpublished*) and nearly half in a South African cohort (19) could not be tested for MMc using this approach due to absence of an informative marker for the mother in the background of the child. In these cohorts, this was secondary to high HLA homology in mothers, as well as a high degree of HLA sharing between mothers and their infants (e.g., non-inherited maternal allele (**NIMA**) matched the paternally inherited allele). In addition, the degree of HLA concordance may influence MMc prevalence and level (19–21), so reliance on HLA-discordance-based assays may exclude key participants and result in biased findings.

Assays that utilize non-HLA global allele frequency data to identify informative markers can also suffer from limitations. Different population groups making up the global frequency data can have vastly different allele frequencies, especially those of African descent. The lack of representation of genomic data sampled from populations other than those of European descent has been noted historically. For example, polymorphism rs2307959 has a minor allele frequency (**MAF**) of 0.4 in European populations but minor allele frequencies of 0 in African and Asian populations in dbSNP. To further illustrate this point, we performed a large query of dbSNP looking for reports of global MAF between 0.40-0.60, but MAF of <0.10 in African populations, and identified 1,129,498 alleles. This demonstrates that global frequency data alone does not predict informativeness in select target populations. Finally, earlier studies have reported chimerism testing using published polymorphisms with relatively high frequencies in the population such as the large deletion polymorphisms in GSTM1 and GSTT1, but both of these variants have functional consequences which could lead to biased data if the alleles are under selective pressure (22).

Given these limitations, we recently developed a low-cost custom approach to detect microchimerism. To increase our ability to identify an informative marker for microchimerism in our populations of interest, we first identified non-HLA polymorphisms with no reported functional impact. We preferentially selected markers that had reported MAF in both African and European descent populations between 0.4-0.6. We further focused on deletion / insertion (**DELINS**) polymorphisms identified in dbSNP and copy number alterations (**CNA**) identified in dbVar and limited our selection to allele targets for which a sensitive and specific qPCR assay could be designed. The resulting approach consists of a multiplex PCR to genotype each sample followed by highly specific companion qPCR assays to assess microchimerism level. Finally, we tested our approach to detect MMc in infant samples from Seattle, Washington, USA and Khayelitsha, Cape Town, South Africa.

## MATERIALS & METHODS

### Selection of polymorphism targets in dbSNP and dbVar

To identify potential polymorphism targets we utilized the filter option in dbSNP (https://www.ncbi.nlm.nih.gov/snp/advanced). Filters were applied with keyword DEL in All Fields, and filters were added for ALFA African population and ALFA European population MAF with range (00000.40-00000.60). The resulting polymorphism list was exported and any variation sequences with less than four base pairs or repeat sequences were removed to increase the ability to create Allele-Specific PCR assays with high specificity. Filtered polymorphisms were exported into Genome Browser (https://genome.ucsc.edu/) and the sequence around the polymorphism was examined. Any with repeated sequences around the DELINS or identified by RepeatMasker as a repeated element were eliminated. For polymorphisms meeting these metrics, the sequence around the polymorphism containing either the DEL or INS sequence was entered into qPCR design software (https://www.idtdna.com/Primerquest/Home/Index). Assays in which allele specific primers could be designed to distinguish the DEL sequence from the INS sequence were moved forward for testing. Among 27 candidates initially identified, both INS and DEL qPCR assays could not be designed for four, DEL qPCR only for 12, and INS qPCR only for one. Thus, 10 candidates were moved forward to multiplex PCR design.

To select for larger DEL sequences, we followed the approach of Aqil, et al (23) to filter for CNA with frequencies between 0.40-0.60 in European and African descent populations in dbVar (https://www.ncbi.nlm.nih.gov/dbvar). CNA sequences with either the DEL or INS sequence were entered into Primer-BLAST (https://www.ncbi.nlm.nih.gov/tools/primer-blast/index.cgi?LINK_LOC=BlastHome). Targets in which Primer-BLAST could design specific primers to both the DEL and INS forms of the polymorphism were selected. The majority of first round potential CNA were eliminated at this stage as specific primers could not be designed for many of the DEL forms of the polymorphism. In addition, Primer-BLAST gave identified homology with other parts of the genome in sequence regions flanking the large INS for many potential targets. Among the initial 20 targets selected, only eight met all metrics and moved to multiplex PCR design.

For both DELINS and CNA candidate polymorphisms, each target sequence was next put into Primer-BLAST and primer sets were configured to create PCR fragments that were compatible with each other and had no target overlap with the goal of creating a multiplex genotyping panel. Selected primer sets were entered into Multiple Primer Analyzer (https://www.thermofisher.com) and primers that showed primer dimer formation with more than two other primers were re-designed. Individual primer sets were then tested on a selection of cell line derived DNA (International Histocompatibility Working Group, Fred Hutch Cancer Center, Seattle USA). PCR fragments representing the DEL and INS forms of each polymorphism were purified using Nucleospin Gel and PCR Clean-up Kit (Macherey-Nagel) and sent for Sanger sequencing (Azenta, South Plainfield USA). Resulting sequence data were analyzed against NCBI reference sequences (https://www.ncbi.nlm.nih.gov/nucleotide/) to confirm the correct target was amplified and corresponded to the expected DEL or INS polymorphism. We additionally adapted two previously published assays (17) to the panel using the same approach, as well as one previously published assay from Olsen, et al (24). For each genotyping target, fluorophores (FAM, SUN, ATTO550, or ATTO565) were added to either the forward or the reverse primer to distinguish individual amplified PCR products for fragment analysis. The set of target assays were then combined into a multiplex PCR to ensure that there was no interference, resulting in the elimination of two DELINS assays, and the final inclusion of 18 assays. Finally, we incorporated a custom PCR targeting amelogenin commonly used to establish XY chromosome status in forensic human identification (25). Primer sequences, fragment size, and allele population frequency information for the final assay set are presented in **Table 1**.

**Table 1.**
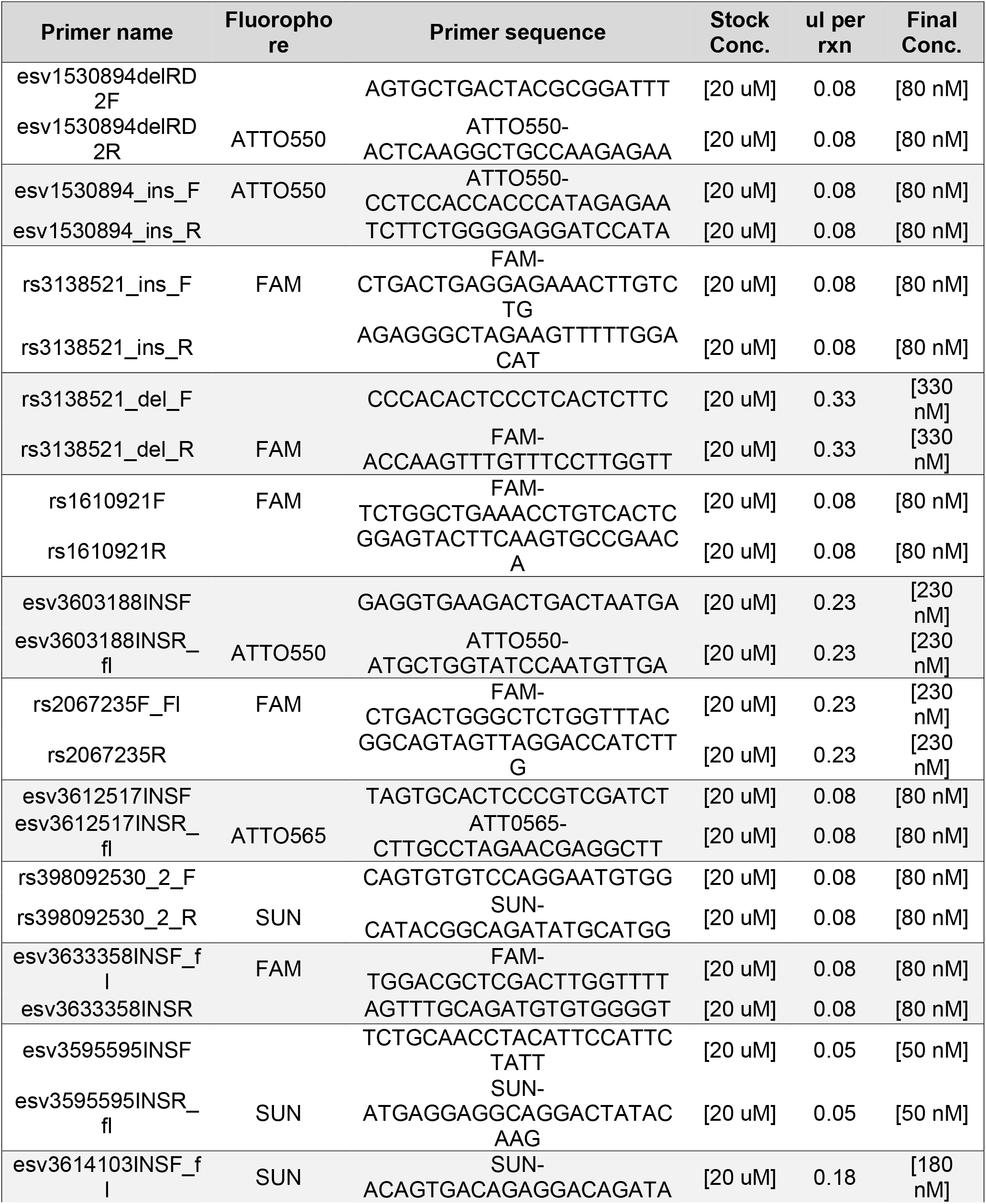
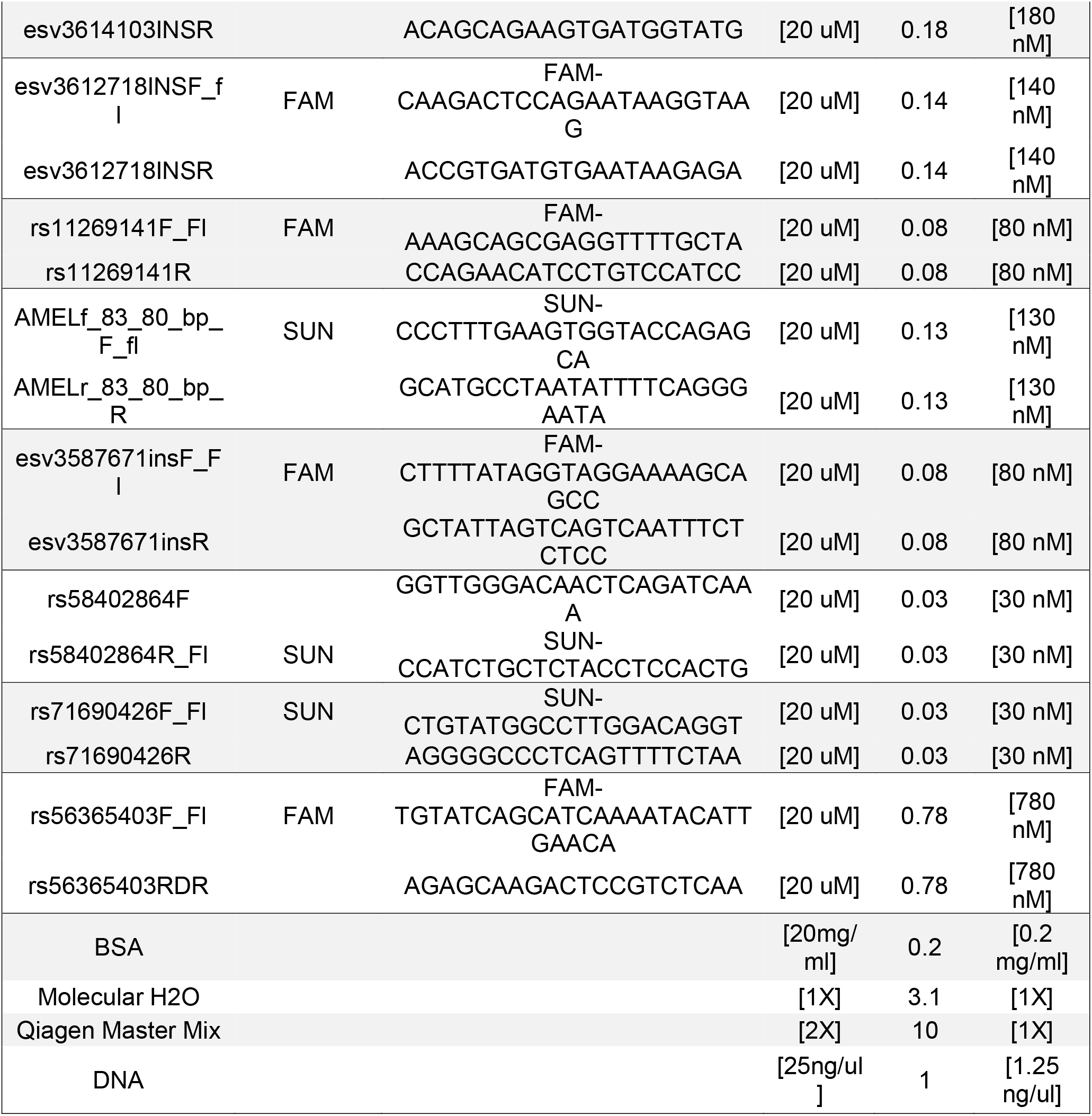
Primer sequences and concentrations for multiplex PCR genotyping reaction.

### Multiplex PCR for genotyping

To identify an informative target for microchimerism, genomic DNA was amplified using the multiplex PCR cocktail (**Table 2**). In brief, a 20 ul PCR reaction contained 10.0 ul of QIAGEN Multiplex PCR Master Mix [2X] (Qiagen), 0.2 ul of Molecular Grade BSA [20 mg/ml], 3.1 ul molecular grade H20, fluorescently labeled and unlabeled primers (idtDNA) of variable amounts, and 25 ng of target genomic DNA. PCR reactions were run in TempAssure PCR 8-tube strips (USA Scientific) on a T100 Thermal Cycler (BioRad) under the following conditions: Taq activation: 95 ° C for 15 min; 40 cycles of Denaturation: 94 ° C for 30 sec, Annealing: 60° C for 1.5 min, and Extension: 72° C for 1 min; and Final Extension: 72° C for 30 min. PCR products (2 ul) were then run on a 4% agarose gel to confirm amplification, followed by dilution 1:30 in molecular grade water and run on a 3730xl DNA Analyzer (ThermoFisher). Resulting .fsa files were imported into the freeware Microsatellite App (Thermofisher) to identify targets present in the donor and not in the recipient for each pair to be assessed (**Figure 1**).

**Table 2.** Expected fragment size on electropherogram and reported allele frequency in dbSNP for each genotyping target.

| African descent allele frequency | European descent allele frequency | Target | Dye label | Expected size (bp) |
| --- | --- | --- | --- | --- |
| 0.586 | 0.475 | esv3587671INS | FAM | 70 |
|  |  | Amelogenin X | SUN | 78 |
|  |  | Amelogenin Y | SUN | 81 |
| 0.858 (38) | 0.282 (38) | rs3138521INS | FAM | 93 |
| 0.142 (38) | 0.718 (38) | rs3138521DEL | FAM | 105 |
| 0.461 | 0.579 | rs146774400DEL | SUN | 106 |
| 0.532 | 0.414 | esv3595595INS | SUN | 118 |
| 0.539 | 0.421 | rs146774400INS | SUN | 130 |
| 0.483 | 0.529 | esv3633358INS | FAM | 132 |
| unknown | 0.560 (39) | esv1530894INS | ATTO 550 | 135 |
| 0.430 | 0.474 | esv3612718INS | FAM | 152 |
| 0.469 | 0.462 | rs1610921DEL | FAM | 161 |
| 0.531 | 0.538 | rs1610921INS | FAM | 169 |
| 0.524 | 0.462 | esv3612517INS | ATTO 565 | 172 |
| 0.477 | 0.543 | rs11269141DEL | FAM | 178 |
| 0.523 | 0.457 | rs11269141INS | FAM | 191 |
| 0.476 | 0.559 | rs58402864DEL | SUN | 198 |
| 0.524 | 0.441 | rs58402864INS | SUN | 211 |
| 0.466 | 0.411 | rs71690426DEL | SUN | 216 |
| 0.546 | 0.649 | rs398092530DEL | SUN | 225 |
| 0.454 | 0.351 | rs398092530INS | SUN | 227 |
| 0.534 | 0.589 | rs71690426INS | SUN | 233 |
| unknown | 0.440 (39) | esv1530894DEL | ATTO 550 | 242 |
| 0.485 | 0.505 | esv3603188INS | ATTO 550 | 255 |
| 0.460 | 0.563 | rs56365403DEL | FAM | 306 |
| 0.540 | 0.437 | rs56365403DEL | FAM | 311 |
| 0.457 | 0.505 | esv3614103INS | SUN | 329 |
| 0.612 | 0.460 | rs2067235DEL | FAM | 356 |
| 0.388 | 0.540 | rs2067235INS | FAM | 366 |

**Figure 1.**
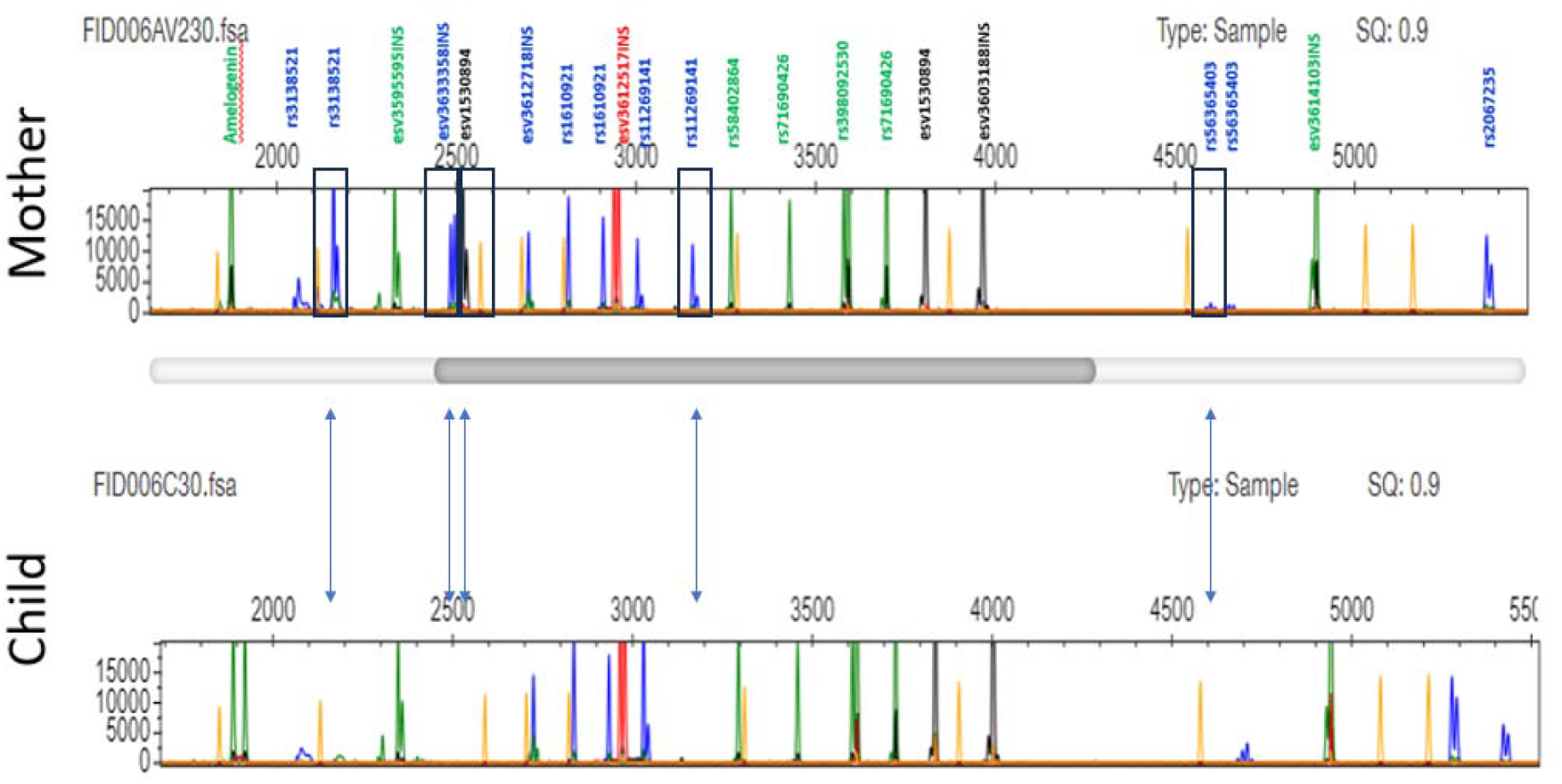
Example of multiplex genotyping electropherograms from a maternal-child pai to identify an informative polymorphism for the detection of maternal microchimerism. Genomic DNA from the mother and her infant were separately assessed using our multiplex PCR genotyping reaction targeting 18 polymorphisms. The PCR product is run on an electropherogram and evaluated using the Microsatellite App (Thermofisher) software. Polymorphisms present in the mother (presence of peak at expected size) and absent in the infant (absence of peak at expected size) are identified as potential targets for the detection of maternal microchimerism. In the selected pair, there are five potential targets highlighted in black boxes in the mother and noted as absent in the child at the position below the blue arrowheads.

### qPCR for microchimerism detection

We next designed a corresponding qPCR assay for each target polymorphism (**Table 3**). Our approach was similar to prior published HLA-based qPCR assays (26), but with the following modifications: the assays were run in a 384 well plate to increase efficiency and sensitivity of fluorescent signal (27, 28), and BSA was added to improve amplification (29, 30). On each 384 well PCR plate two qPCR assays were run in parallel: one for the target of interest and the other for betaglobin to assess genomic input of the recipient DNA. Standard curves for each assay were run on the same plate for quantitation, and wells representing 2 and 0.5 genomic equivalent (**GE**) of the polymorphisms assay were run to assess assay sensitivity. Due to the high sensitivity of the assay, all PCR set-up was performed in a laminar hood with UV sterilization capabilities (AirClean 600 PCR Workstation, AirClean Systems, Creedmoor, USA), and all surfaces and tools were cleaned prior to each plate setup using 10% bleach followed by 70% ethanol.

**Table 3.** qPCR primer and probe sequences for assessment of microchimerism.

The standard curve for each polymorphism target was created from a source of renewable genomic DNA (International Histocompatibility Working Group cell lines, Fred Hutch Cancer Center, Seattle, USA). The standard curves for each polymorphism target included the following points to ensure linearity and to confirm sensitivity: 2,048GE, 512GE, 128GE, 32GE, 8GE, 2GE, 0.5GE, and 0GE (no template control) and were run in triplicate (**Figure 2**). In two of the three polymorphism standard curves 8,000 GE (5 ul of 1,600GE/ul) of cell line genomic DNA negative for the polymorphism being targeted was added to each well to simulate background off-target genomic DNA. The standard curve for the betaglobin assay was generated using commercially available human genomic DNA (Human Genomic DNA, Promega, Madison, USA) and included the following points: 40,960GE, 10240GE, 2560GE, 640GE, 160GE, 40GE, 10GE, and 0GE (no template control). All standard curves were created in a 10mM Tris-Cl buffer pH 8.5. Recipient DNA to be tested for microchimerism was diluted to approximately ∼1,600 GE/ul in a total volume of 105 ul. Five microliters of recipient DNA was added to each of 18 wells (polymorphism assay) and 2.5 ul of recipient DNA was added to 2 wells (betaglobin assay). Molecular grade water was added to bring the contents of each well to 10 ul, followed by the addition of 15 ul of qPCR cocktail to bring the final volume to 25 ul. The qPCR cocktail included [1X] TaqMan® Universal PCR Master Mix (Thermo Fisher Scientific, Carlsbad, USA), [300nM] each primer, [100nM] of probe, and [0.2ug/ul] Molecular Grade BSA (Thermo Fisher Scientific, Waltham, USA) for all assays excluding betaglobin, esv1530894, and rs3138521.

**Figure 2.**
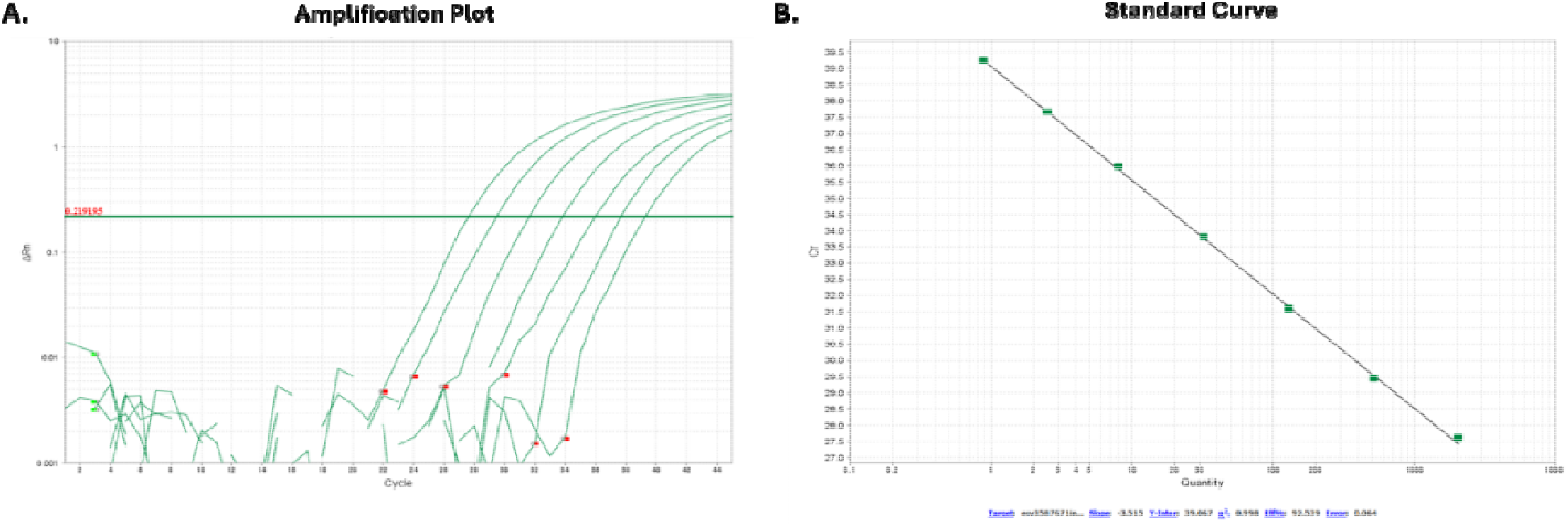
Example of qPCR assay sensitivity for one of the custom designed targets (esv3587671ins). A) Amplification curves for standard curve points 2,048GE, 512GE, 128GE, 32GE, and 8GE and positive control wells 2GE and 0.5GE. B) Standard curve points for 2,048GE, 512GE, 128GE, 32GE, and 8GE and positive control wells 2GE and 0.5GE plotted on a best fit line. Points demonstrate excellent linearity with R^2^=0.998 and expected slope (-3.5).

For each assay, the plate was cycled in a QuantStudio 5 (Thermo Fisher Scientific, Carlsbad USA) under the following conditions: 50°C for 2 min (Uracyl-N-glycosylase to eliminate potential PCR product contamination); 95°C for 10 min (Taq activation); and 45 cycles of 95°C (Denaturation) for 15 sec and 60°C (Annealing) for 1 min. Data collection was set at the 60°C (Annealing) step. After each run, assay data was analyzed in the QuantStudio™ Design and Analysis Software. Quality control (**QC**) of the polymorphism standard curve was performed: coefficient of determination (R^2^) greater than 0.955, slope -3.3 to -3.7 (optimal -3.5), amplification of at least 2 of the duplicate wells of 0.5GE or 2GE points (sensitivity), and all 0GE wells devoid of amplification (specificity). The best fit line for standard curve included all points above 2GE, whereas 0.5GE and 2GE were used only as positive controls. Polymorphism standard curves and any positive test wells were normalized to the betaglobin standard curve. These normalized test well values were then summed and the level of microchimerism in the recipient DNA was quantified as total polymorphism specific GE over total betaglobin GE tested.

Over multiple runs for each polymorphism assay, the standard curves for were assessed for sensitivity and reproducibility. All standard curve runs for all assays had R^2^ of 0.99 or greater. The average slope was -3.42 (-3.27 to -3.73), and in all runs the positive control points 0.5GE and / or 2GE amplified in at least two wells. However, the 0.5GE point did not amplify in duplicate in all runs so calculation of reproducible lowest limit of sensitivity was considered to be 2GE for the assays. This pattern was seen in previous qPCR runs targeting HLA-based assays performed in our laboratory (data not shown). The average %CV of the 2GE duplicates over all runs was 1.92% (0.03%-5.72%). The 2GE point represents 2GE input into an off-target background of 8,000GE, indicating a reproducible sensitivity limit of these assays of 0.025%.

### Extension of the approach to nested PCR to further reduce cost

To further reduce costs, we developed a related nested PCR based approach for detection of microchimerism that utilizes a single qPCR well to generate a semi-quantitative result. Details can be found in the **Supplement**.

### Cohorts

We tested our method in three cohorts. The first was the Fetal Immune Development (**FID**) study based in Seattle, WA, USA (31). The cohort enrolled women with singleton pregnancies receiving obstetric care at the University of Washington affiliated clinics at the time of their first prenatal visit and followed them across pregnancy. Peripheral blood mononuclear cells (**PBMC**) were isolated from maternal whole blood collected into ACD tubes at the time of delivery. Cord blood was collected into ACD tubes via cross-clamping, cleaning, and direct cannulation of the cord, and cord blood mononuclear cells (**CBMC**) were isolated. The study was approved by the UW Institutional Review Board (**IRB**) (STUDY00001636, STUDY00004878). The second was the Innate Factors Associated with Nursing Transmission (**InFANT**) Study based in Khayelitsha, Cape Town, South Africa, which enrolled mothers and their infants at delivery, including mother living both with and without HIV (19). Maternal blood was collected at delivery and whole blood was collected from infants one day one of life (“birth”) and fixed 1:10 with 1X FACS Lysing Buffer (BD Biosciences), pellet, washed, and cryopreserved. The study was approved by University of Cape Town’s Human Research Ethics Committee (Rec 285/2012) and Seattle Children’s IRB (Protocol 15690). The third was the Maternal immunizations in low-and high-risk pregnancies (**MATIMM**) study (**Supplement**), which enrolled women and their infants with blood collected at delivery and during infancy and processed for PBMC. The study was approved by the University of Washington IRB (STUDY00008491). All women provided written informed consent for themselves and their infants in all three cohorts.

## RESULTS

### Identification of maternal microchimerism in two cohorts

We next tested the method to detect MMc in two cohorts, the FID cohort from the United States for 69 pairs and the InFANT cohort from South Africa for 125 pairs. For the Seattle cohort, our tested substate included T cells and non-T cells sorted from CBMC by magnetic activated cell sorting (MACS). Briefly, MACS sorting was performed using a CD3 Ab (Biolegend, San Diego, USA) and MS columns (Miltenyi Biotec, Gaithersburg, USA) following manufacturer’s protocol. For the Cape Town cohort our substrate was fixed whole blood from birth. DNA was extracted from MACs sorted T cells and birth fixed whole blood from the InFANT Study with a modification of the PrepFiler™ Forensic DNA Extraction Kit (Thermo Fisher Scientific, Carlsbad, USA) to include spike-in of 1 ug of carrier RNA (Qiagen, Germantown, USA) and two sequential elutions to increase yield. The non-T cells isolated by MACS were extracted using PureLink™ DNA Mini Kit (Thermofisher Scientific) following the manufacturer’s protocol.

Within both cohorts, an informative marker for MMc was identified by the multiplex genotyping assay in all pairs. For the Seattle cohort, 16 pairs (24%) had one informative marker for MMc, 22 (32%) had two, 15 (22%) had three, nine (13%) had four, two (3%) had five, two (3%) had six, one (2%) had seven, and two (3%) had eight. In the Cape Town population, 29 pairs (25%) had one informative marker for MMc, 28 (24%) had two, 27 (23%) had three, 13 (11%) had four, 14 (12%) had five, two (2%) had six, three (3%) had seven, and two (2%) had eight. In cases in which more than one marker was found, samples were triaged into groups with the same informative marker for efficiency in qPCR screening. We identified MMc in 47 of 68 (69%) T cell fractions and 33 of 68 (49%) non-T cell fractions from the FID cohort (**Table 4**). Level of MMc amongst positive T cell fractions ranged from 0.5-3,088 per 10^5^ infant GE and amongst non-T cell fractions 0.6-2,223 per 10^5^ infant GE. Within birth fixed whole blood from the InFANT cohort, 96 of 118 (81%) samples were positive for MMc with levels that ranged from 0.4-419 per 10^5^ infant GE (**Table 4**). For each of the three sample sets (FID T cells, FID non-T cells, InFANT birth blood), the GE tested in samples that tested MMc negative vs. MMc positive was not statistically different.

**Table 4.** Prevalence and level of MMc in United States and South African Cohorts.

| Cohort | Total (n) | MMc negative |  | MMc positive |  |  |
| --- | --- | --- | --- | --- | --- | --- |
|  |  | n (%) | Genomic Equivalents | n (%) | Genomic Equivalents | MMc Level |
| FID T cells | 68 | 21 (31%) | 91,148<br>(6,360-194,127) | 47<br>(69%) | 83,787<br>(4,385-232,893) | 4 (0.5-3,088) |
| FID Non-T cells | 68 | 35 (52%) | 108,502<br>(4,907-166,975) | 33<br>(48%) | 93,718<br>(2,073-198,142) | 3 (0.6-2,223) |
| InFANT whole birth blood | 118 | 22 (19%) | 148,379<br>(15,676-223,642) | 96<br>(81%) | 142,342<br>(3,276-229,957) | 3 (0.4-419) |

*Genomic equivalents of recipient tested, presented as median (range). MMc level expressed per 100,000, presented as median (range) of positive samples.*

We next explored if assessment of MMc using two different informative targets would yield concordant findings. A subset (n=8) of samples from the Cape Town cohort had more than one target polymorphism for MMc identified, were positive for MMc on initial testing, and had sufficient DNA for repeat testing. Overall, we saw some variation in absolute MMc level detected but all samples had MMc detected in both runs (**Table 5**). Further, as assessed with paired t-tests, there was no statistical difference in the number of positive wells (p=0.1) or MMc level identified (p=0.5) between the first and second assay for each sample, indicating that different markers yielded similar findings.

**Table 5.**
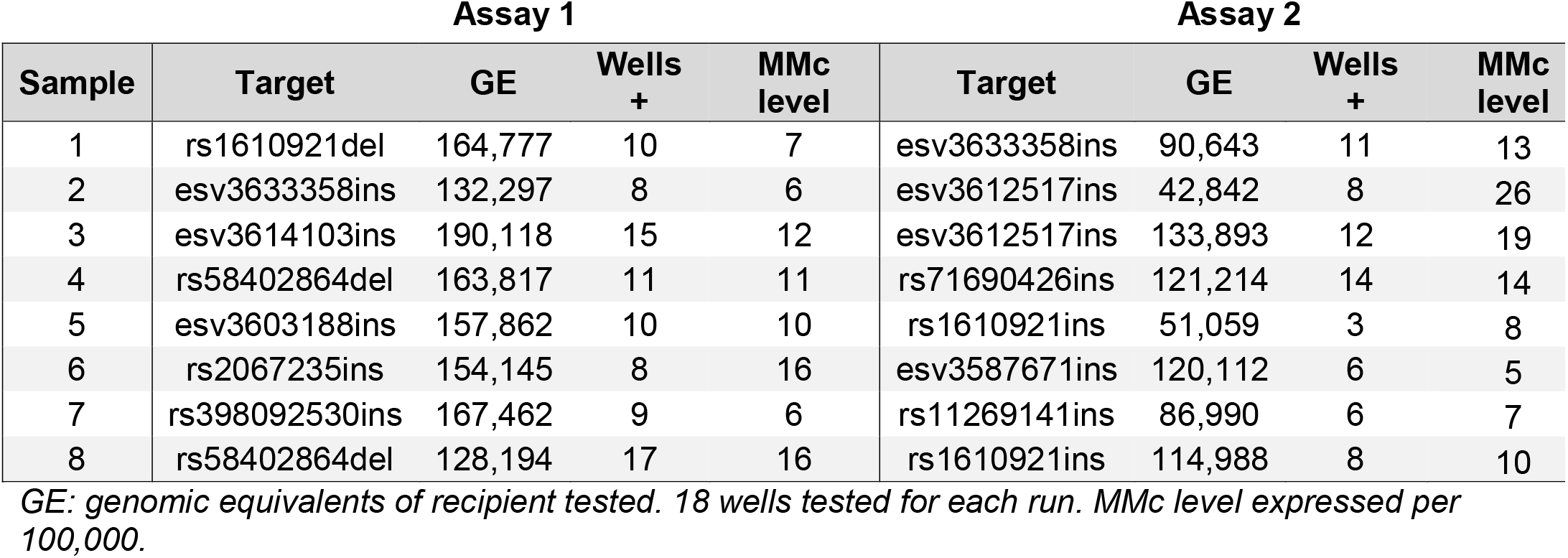
Select testing of South African samples with two informative targets for MMc.

| Assay 1 |  |  |  |  | Assay 2 |  |  |  |
| --- | --- | --- | --- | --- | --- | --- | --- | --- |
| Sample | Target | GE | Wells<br>+ | MMc<br>level | Target | GE | Wells<br>+ | MMc<br>level |
| 1 | rs1610921del | 164,777 | 10 | 7 | esv3633358ins | 90,643 | 11 | 13 |
| 2 | esv3633358ins | 132,297 | 8 | 6 | esv3612517ins | 42,842 | 8 | 26 |
| 3 | esv3614103ins | 190,118 | 15 | 12 | esv3612517ins | 133,893 | 12 | 19 |
| 4 | rs58402864del | 163,817 | 11 | 11 | rs71690426ins | 121,214 | 14 | 14 |
| 5 | esv3603188ins | 157,862 | 10 | 10 | rs1610921ins | 51,059 | 3 | 8 |
| 6 | rs2067235ins | 154,145 | 8 | 16 | esv3587671ins | 120,112 | 6 | 5 |
| 7 | rs398092530ins | 167,462 | 9 | 6 | rs11269141ins | 86,990 | 6 | 7 |
| 8 | rs58402864del | 128,194 | 17 | 16 | rs1610921ins | 114,988 | 8 | 10 |

*GE: genomic equivalents of recipient tested. 18 wells tested for each run. MMc level expressed per 100,000.*

## DISCUSSION

Herein we present a set of novel qPCR assays for the detection of microchimerism as an alternative to existing assays, many of which rely on polymorphisms in the HLA Class II region. These assays circumvent the high cost of HLA typing and the lack of informative HLA-based markers identified in our studies of maternal-child pairs of African descent (19, 32, 33). Current non-HLA based approaches for the detection of chimerism were either limiting in cost or sensitivity or did not have markers that were suitable for populations of African descent. As such, we were motivated to assess a wider pool of polymorphisms, considering both DELINs and CNA from dbSNP and dbVar, and pre-selecting markers with high allele prevalence in our populations of interest. We subsequently based the design of our qPCR assays targeting these alleles on prior rigorous assays published in the field (17, 18) but added the novel features of a multiplex-PCR based approach for genotyping and translation of the assays to a higher throughput format to increase efficiency. Our subsequent panel of qPCR-based assays for the detection of microchimerism are low cost, highly sensitive and specific, and designed to include targets for a wide variety of populations, including both those of African and European descent.

We modeled our multiplex PCR genotyping assay on the Qiagen Investigator assay (34) and deliberately incorporated the forensic human identification target amelogenin targeting the XY chromosomes as an additional QC to confirm sample identity. Successful amplification of multiple PCR targets in one assay, like in the Qiagen assay, was challenging and required primer titration and re-design of the panel multiple times. Interestingly, in early development attempts, we tried commercially available multiplex design software but found that they did not accurately reflect bench performance. We also tried the use of PCR additives such as betaine and Q solution but found they varied in impact to each individual target rather than leading to overall improvement in assay performance.

In assessing our companion qPCR assays, we determined that our assays match the sensitivity and specificity of previously published assays (17, 18). We note that a similar approach by Olsen, et al (24) was recently published, and we attempted to incorporate a subset of their qPCR assays into our own existing panel. Amongst the ten assays, four were chosen that had reasonable frequencies in populations of African descent. However, in our hands, we were not able to achieve the specificity of our other qPCR assays; in particular, we identified off-target amplification for assays targeting the deletion form of the polymorphisms. We retained one target (rs398092530) but re-designed the deletion assay and created a blocking primer to suppress off target amplification. Finally, our development testing indicated in conditions of low substrate concentration or low microchimerism levels, the assay was robust using a fixed number of input wells, eliminating the need for gDNA concentration and facilitating throughput (data not shown).

We assessed our method in two maternal-infant cohorts from Seattle, USA and Cape Town, South Africa and identified a targetable polymorphism for the detection of MMc in all infants. This represents a significant improvement over prior HLA-based approaches, in which a large proportion of pairs were excluded from testing based on the inability to identify a unique marker of the mother in the background of the infant. Further, 76% of pairs from Seattle and 75% of pairs from Cape Town had more than one polymorphism that could be targeted, allowing for the selection of a target shared with other samples and thereby increasing efficiency. Using our companion qPCR assays, we identified T cell MMc in 69% of infant cord bloods from Seattle and non-T cell MMc in 48%. In South Africa, we identified MMc in 81% of fixed whole blood from birth. These proportions are consistent with the range of MMc reported in other birth samples, as were the MMc levels we identified (17, 19, 32, 33). Finally, in a subset of individuals from South Africa in whom we could directly compare the performance of two informative markers, there was good concordance between the assays. These data indicate that our approach allows for robust characterization of microchimerism in populations of variable regional and ethnic descent.

Our approach has some limitations. One challenge in the field in general is the stochastic nature of detection of low level microchimerism, in which as low as one copy of an allele is targeted in a high concentration of off-target background. We identified a similar finding in the reproducibility of our very low input positive controls (0.5GE), and thus we have chosen to report the reproducible limit of sensitivity of the assays based on the 2GE positive control (0.025%). Indeed, others have reported differences in microchimerism detection in testing of the same sample multiple times, reflecting this stochastic distribution (24, 35). Further, although we identified complete “informativeness” in our two cohorts from Seattle and Cape Town, this may not be reflective of other cohorts. Future work should assess performance of the method in other populations of African and non-African descent with refinement of polymorphism selection as needed. Finally, we explored moving our assays from a traditional qPCR-based approach to more modern digital PCR, however, we identified some loss of sensitivity at very low levels of microchimerism (data not shown). Further extension of the assay to newly developed platforms such as Ultra-PCR (36) or Solid-Phase PCR (37) could potentially increase throughput or sensitivity.

In conclusion, we present a panel of non-HLA polymorphism assays for the sensitive and specific detection of microchimerism. Our method decreases cost while maintaining rigorous standards and, perhaps most importantly, empowers the study of microchimerism in diverse human populations.

## ACKNOWLEDGEMENTS

We would like to acknowledge the assistance of Cara Tobey and Jia Wong in recruiting participants and the assistance of Quinn Peters, Abigail Spray, Aiesha Abdulla, and Nolawit Mulugeta in processing samples from the FID cohort.

