## Supplemental Data for "A sensitive and specific microchimerism detection method using custom designed genotyping and companion qPCR assays"

**Supplemental Material**

**Adaptation of qPCR assays to a nested PCR approach to increase efficiency**

Our current protocol assesses samples for MMc level across 18 qPCR wells. We explored whether we could increase efficiency by developing a nested PCR reaction as an alternative. We took advantage of the CNA polymorphisms in our panel to design nested PCR reactions. The CNA polymorphisms are large DELINS polymorphisms greater than 100 bp, which allows assay design room to create nested PCR assays with specificity and sensitivity. For the targets esv3587671ins, esv3595595ins, esv3633358ins, esv3612718ins, esv3612517ins, esv3603188ins, and esv3614103ins nested PCR primers were designed flanking either side of the qPCR primers (**Table S1**). For input we used 100 ng of test DNA (~15,150 GE) in the primary PCR reaction in a cocktail consisting of [1X] QIAGEN Multiplex PCR Master Mix (Qiagen), [400nM] each primary nest primer, [1X] Q solution, and [1X] Molecular Grade H2O. The PCR was run in 384 well GeneAmp® PCR System 9700 (Thermo Fisher Scientific) under the following conditions: Taq activation: 95 ° C for 15 min; 40 cycles of Denaturation: 95 ° C for 30 sec, Annealing: 60° C for 1 min, and Extension: 72° C for 1 min; and Final Extension: 72° C for 5 min. For the secondary PCR reaction, 2 ul of the primary PCR reaction was assayed in a single well with the MMc specific qPCR assay using the target primers as in our standard protocol.

We tested this approach to assess for MMc in granulocytes in infants from the MATIMM cohort. In brief, blood samples collected from infants were processed using Lymphocyte Separation Medium (Corning, Manassas, USA) for isolation of PBMCs following manufacturer’s protocol. Post collection of PBMCs, the remaining fraction containing red blood cells and granulocytes was further treated for cell isolation. Remaining Lymphocyte Separation Medium was removed by pipet leaving the cell layers at the bottom of the tube. The sample was diluted with Hank’s Balanced Salt Solution [1X] (Thermo Fisher Scientific) and centrifuged at room temperature at 300 g for 5 minutes with brake set at 5. The clear supernatant was removed by pipet leaving the remaining cells and red blood cells. RBC Lysis Buffer [1X] (Biolegend, San Diego, USA) was added and tube was inverted to mix and allowed to incubate for 20 minutes. The sample was centrifuged at room temperature at 300 g for 5 minutes with brake set at 5. Lysed red cells were removed by pipet leaving the cell pellet. The pellet was resuspended in PBS and transferred to microcentrifuge tubes for gDNA extraction. DNA was extracted from the granulocyte cell pellets using the PureLink™ DNA Mini Kit (Thermofisher Scientific) following the manufacturer’s protocol. Ten samples in which the MMc target was one of the seven CNA polymorphisms were tested using our nested PCR screening protocol. Samples were categorized as positive or negative for MMc according to amplification in the qPCR reaction in a single well that utilized the nested PCR product as input. To confirm our nested MMc screen correctly identified MMc positive samples, we tested the same sample using our standard 18 well qPCR protocol. The nested assay correctly identified the samples in which MMc was detected at a wide range of levels (**Table S2**).

**Supplemental Table 1. Nested PCR primers**

| **Primer name** | **Primer sequence** |
| --- | --- |
| esv3603188_TqMn_F | CTGTCTGGCTCTCTTTCTCTTG |
| esv3603188INSTQNSTR | ACAGACTATCACAGACTGGGT |
| esv3587671NstF1 | GCTTTTATAGGTAGGAAAAGCAGCC |
| esv3587671NstR1 | TTGATAGACTCTTAAAGGCAGCTT |
| esv3612517INSTQNSTF | AGGCTCCTGGGTTCCTATGT |
| esv3612517INSTQNSTR | GCCATGGAATGATGTGTTGA |
| esv3633358INSTQNSTF | CCTGCTCAAACAAAATGTGC |
| esv3633358INSTQNSTR | TCAAAGGAGAGTGGGATGGT |
| esv3595595INSF | TCTGCAACCTACATTCCATTCTATT |
| esv3595595INSR | ATGAGGAGGCAGGACTATACAAG |
| esv3614103INSF | ACAGTGACAGAGGACAGATA |
| esv3614103INSR | ACAGCAGAAGTGATGGTATG |
| esv3612718INSF | CAAGACTCCAGAATAAGGTAAG |
| esv3612718INSR | ACCGTGATGTGAATAAGAGA |

**Supplemental Table 2. Concordance of qPCR and nested PCR for detection of MMc**

| **Sample** | **Assay** | **GE assessed** | **Wells + (n)** | **MMc Level** | **Nested PCR Result** |
| --- | --- | --- | --- | --- | --- |
| 1 | esv3633358INS | 176,633 | 3 | 2 | MMc+ |
| 2 | esv3633358INS | 152,863 | 18 | 62 | MMc+ |
| 3 | esv3612517INS | 88,615 | 18 | 360 | MMc+ |
| 4 | esv3603188INS | 196,452 | 16 | 20 | MMc+ |
| 5 | esv3612517INS | 150,021 | 0 | 0 | MMc- |
| 6 | esv3603188INS | 163,329 | 1 | 1 | MMc+ |
| 7 | esv3633358INS | 177,147 | 0 | 0 | MMc- |
| 8 | esv3633358INS | 156,151 | 0 | 0 | MMc- |
| 9 | esv3612517INS | 144,773 | 0 | 0 | MMc- |
| 10 | esv3603188INS | 174,306 | 7 | 5 | MMc+ |
| *All samples tested by qPCR across 18 wells, and via a single well for the nested PCR approach. MMc level expressed per 100,000 GE tested.* | | | | | |
